# Many options, few solutions: over 60 million years snakes converged on a few optimal venom formulations

**DOI:** 10.1101/459073

**Authors:** Agneesh Barua, Alexander S. Mikheyev

## Abstract

Gene expression changes contribute to complex trait variations in both individuals and populations. However, how gene expression influences changes of complex traits over macroevolutionary timescales remains poorly understood. Being comprised of proteinaceous cocktails, snake venoms are unique in that the expression of each toxin can be quantified and mapped to a distinct genomic locus and traced for millions of years. Using a phylogenetic generalized linear mixed model, we analysed expression data of toxin genes from 52 snake species spanning the three venomous snake families, and estimated phylogenetic covariance, which acts as a measure of evolutionary constraint. We find that evolution of toxin combinations is not constrained. However, while all combinations are in principle possible, the actual dimensionality of phylomorphic space is low, with envenomation strategies focused around only four major toxins: metalloproteases, three-finger toxins, serine proteases, and phospholipases A2. While most extant snakes prioritize either a single or a combination of major toxins, they are repeatedly recruited and lost. We find that over macroevolutionary timescales the venom phenotypes were not shaped by phylogenetic constraints, which include important microevolutionary constraints such as epistasis and pleiotropy, but more likely by ecological filtering that permits a few optimal solutions. As a result, phenotypic optima were repeatedly attained by distantly related species. These results indicate that venoms evolve by selection on biochemistry of prey envenomation, which permit diversity though parallelism and impose strong limits, since only a few of the theoretically possible strategies seem to work well and are observed in extant snakes.

## Introduction

Single genes underlying major traits are the exception rather than the rule, and the dissection of polygenic trait variation has been at the forefront of biological research (1–3). Much of the complexity resulting from interactions between genes is mediated through their expression, which plays a central role in determining phenotypic variation between individuals and populations (4–8). In particular, levels of gene expression account for substantial sources of variation in natural populations, acting as potential targets of natural selection (8–10). Although population-level differences in expression may contribute to the onset of local adaptation and perhaps even eventual adaptive divergence (6, 11, 12), how changes in gene expression levels lead to evolution of complex traits over the course of millions of years remains largely unknown.

Interactions between genes and their effect in channelling of adaptive responses have been the focus of the field of quantitative genetics. How evolution results from the combined effects of the adaptive landscape, and the pattern of genetic variances and covariance among genes (the G matrix), is one of the key questions in this field (13, 14). The covariance between genes plays a vital role in shaping complex traits by determining the evolutionary trajectory through natural selection (15), and the occurrence of parallelism (16). While most quantitative genetics studies deal with populations, their conclusions can translate to macroevolutionary processes as well. For example, estimates of divergence between populations show that the direction of greatest phenotypic divergence can be predicted by the multivariate direction of greatest additive genetic variance within populations (17). Unfortunately, the G matrix cannot be extrapolated across macroevolutionary timescales, as it itself evolves (18). Fortunately, it is possible to compute a phylogenetic covariance matrix for multivariate traits, which can serve as a useful analogy to the G matrix, but over much larger timescales, and incorporating a broader range of constraints (19, 20). We can then examine whether the structure of the phylogenetic covariance matrix corresponds to evolutionary trajectories of complex traits.

Here we use the analogy between the G matrix and the phylogenetic covariance matrix to understand how gene expression evolves in a complex trait, namely snake venom. Being composed of proteinaceous cocktails, snake venoms are unique in that the expression of each toxin type can be quantified and traced to a distinct genomic locus (21–23). Variations in gene expression alter the abundance of proteins in the venom, thereby influencing venom efficacy (24–26). Thus, toxin expression levels constitute the polygenic phenotype that is the venom, allowing us to examine how selection affects gene expression over tens of millions of years. To examine the features of complex trait evolution at the level of gene expression, we estimated phylogenetic covariance of 10 toxins using data from 52 snake species covering the three venomous snake families (Elapidae, Viperidae, and Colubridae) and asked the extent to which it matched observed patterns of evolutionary change across taxa.

Although we find that extant snake venoms occupy a limited area of phenotypic space, largely centred around four major toxin families, there are no phylogenetic constraints to the number of possible venom combinations. These data show that the relatively small number of molecular strategies used by the snakes result from consistent and often convergent selection on the biochemistry of envenomation, rather than from intrinsic constraints on gene interactions. Thus, over tens of millions of years selection likely plays a greater role in shaping the venom phenotype than intrinsic constraints.

## Results

### Expression data and phylogeny

Expression data for snakes were collected from published studies that reported relative levels of toxin expression via next-generation (Illumina and 454) transcriptome sequencing of cDNA libraries. We obtained data for a total of 52 different snake species from the three major venomous families (Colubridae, Elapidae and Viperidae), from a list of 39 publications (Supplementary Table 1). For inclusion, each study had to provide quantitative data on toxin component abundance and had species for which phylogenetic data were available. We restricted our dataset to include components that are found in at least 50% of snakes to focus on generally important toxins, and because sample sizes for the other components would be too low for accurate and phylogenetically unbiased inference. Overall 10 out of 27 toxins we retained. For comparative analyses, we used a published time-calibrated phylogeny of squamates, which estimated the most recent common ancestor (root) of the three snake families to about 60 million years ago (27).

### Evolutionary covariance between venom components

By limiting the range of responses to natural selection, the covariances between genes reflects constraints that shape a phenotype. The phylogenetic covariance matrix (PCOV) accounts for the effect of phylogeny on the interrelationships between genes coding for the snake venom phenotype, providing an approximation of the presence or absence of constraint behind the evolution of gene expression levels. To estimate the PCOV, we used a phylogenetic generalized linear mixed model (PGLMM) under a Bayesian framework. The concept of PGLMM was devised in the early 90s as a method to infer evolutionary constraints of characters using only phylogeny and measures of phenotypes (19). As an extension of maximum likelihood based techniques widely used in quantitative genetics, PGLMM was notable for its versatility as a comparative method (28, 29). We use a modern rendition of the PGLMM devised by Hadfield and Nakagawa, which was optimized for faster and better performance (29, 30). The PGLMM estimated changes in gene expression against a presumed change in diet with the effect of phylogeny being modelled as a random effect. Life history characteristics and diets for snakes are difficult to obtain, particularly in a consistent manner. However, a snake’s potential diet is largely affected by its body size, with smaller individuals consuming smaller prey, while larger individuals tend to prefer larger prey (31). Therefore, we used adult snake length as a proxy for diet. The mean effective sample size for all parameters was greater than 11,000 (Supplementary figure 4). The diagnostics revealed suitable convergence of the chains with negligible autocorrelation in the MCMC (Supplementary Fig. 1-3). Significant values in the PCOV matrix denote the presence of phylogenetic constraint, while non-significant values denote its absence. We observed a lack of significant values in the PCOV (Fig. 1) for all the venom components that we modelled. In addition to estimating a PCOV, the model was used to compute λ values which denote the phylogenetic signal (Fig. 1), similar to Pagel’s λ model for phylogenetic signal (29). The λ values are a measure of statistical dependence of trait values and phylogeny. They indicate whether certain components in modern snakes were likely similar as in their ancestors. In our case, most venom components show strong phylogenetic signals of greater that 0.5, albeit with large confidence intervals. However, and all venom components have λ significantly greater than zero. A few, in particular cysteine-rich secretory proteins (CRISPs), metalloproteinase (SVMP), three finger toxin (TFTx), and Kunitz-type serine protease inhibitor (KSPI) show very strong phylogenetic signals (> 0.8) and narrow confidence intervals, indicating the presence of strong phylogenetic inertia.

**Fig. 1.**
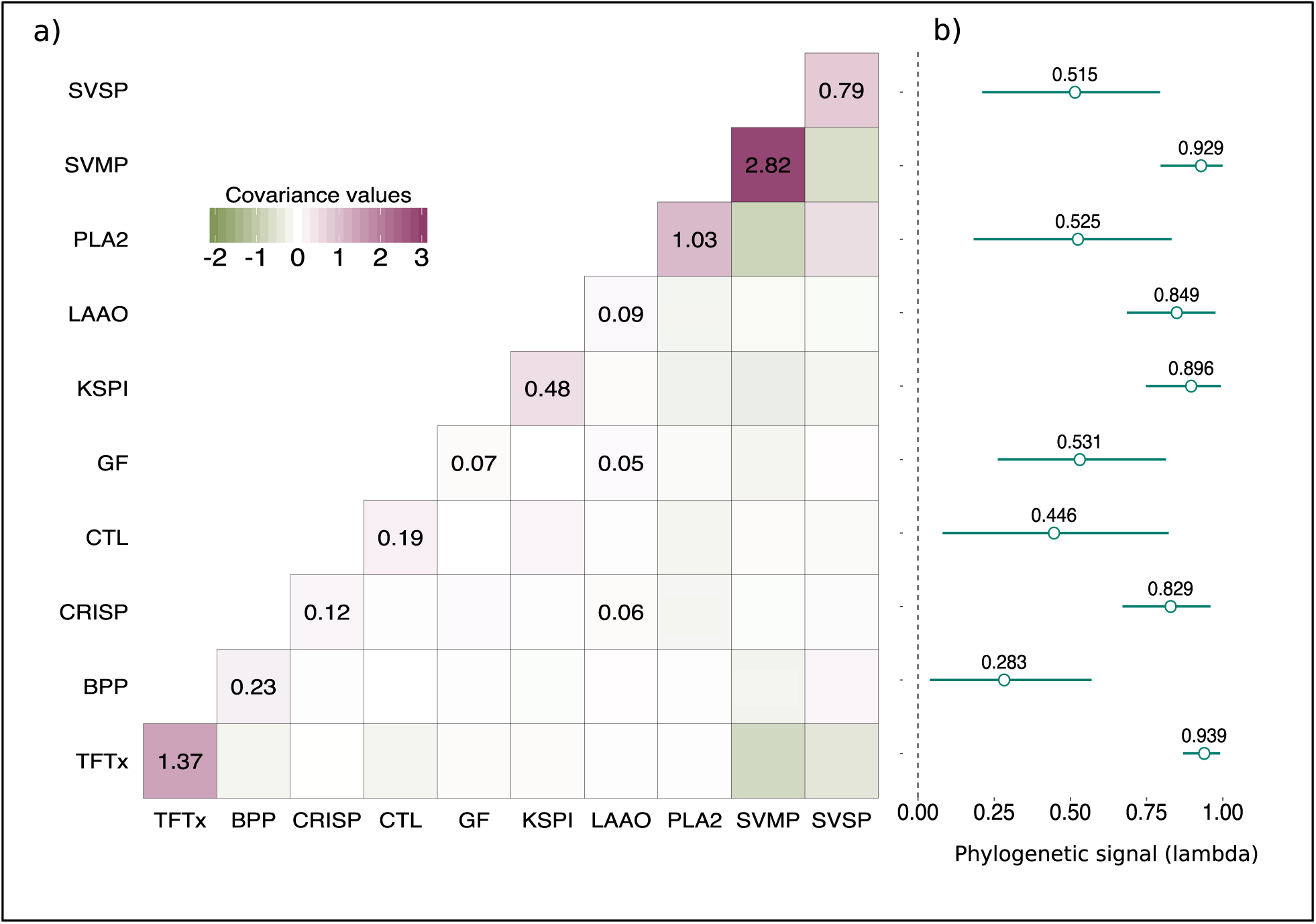
Phylogenetic constraints on individual toxins and their combinations. **a**, A lack of significant values (only significant values labelled) in the phylogenetic covariance matrix denote a lack of phylogenetic constraint between toxins. **b**, Components show a significant presence of a phylogenetic signal, indicating that closer species are likely to evolve the same way. Lambda, represents phylogenetic signal, which is a measure of dependency of trait evolution with phylogeny. Lambda values, are estimated as toxin variance on the diagonal, divided by the sum of diagonal variance and residuals. TFTx, SVMP, KSPI, LAAO, and CRISP showed the highest signal, with greatest significance, while the rest showed comparatively weaker signals. Phylogenetic constraints determine convergence and parallel evolution, where high constraint reduces the likelihood of genes contributing to different convergent regimes (16). Yet, for snake venom genes we see no such constraints in gene expression despite the high phylogenetic signal, suggesting that all toxin combinations, in principle, are possible.

### Four toxins drive the evolution of the snake venom arsenal

We identified axes of maximum variations in the toxin components using PCA on the phylogenetic covariances, using it to visualize the dimensionality of the venom phenotype (32). The first two components, which jointly explained 73.6% of the variation, had the largest loadings from four families of toxins: three finger toxins (TFTx), snake venom metalloproteinase (SVMP), phospholipase A2 (PLA2), and snake venom serine protease (SVSP) (Fig. 2). We therefore classified them as ‘major’ toxins, representing three largely distinct envenomation strategies focussed around SVMP, TFTx, and a combination of PLA2 and SVSP.

**Fig. 2:**
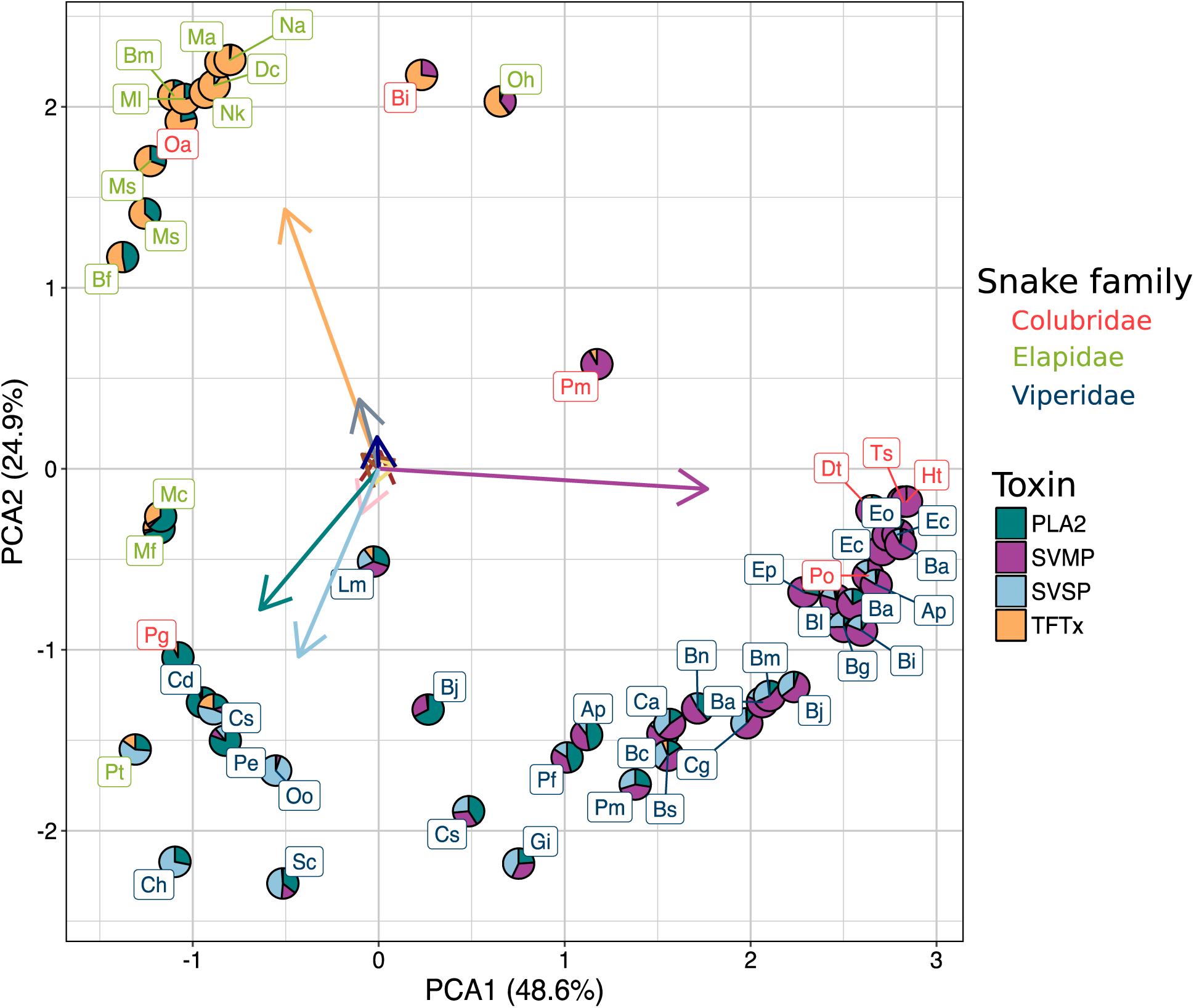
Snakes cluster on phylomorphospace along the axes of four toxins: Phospholipase A2 (PLA2), Snake venom serine proteases (SVSP), snake venom metalloproteases (SVMP), and three finger toxins (TFTx), (Species codes in Supplementary Note 1). These axes represent three distinct envenomation strategies employed by the snakes. Vipers in our data employ a wide spectrum of strategies, from being focussed primarily on SVMP, to employing a mixture of PLA2 and SVSP. Most elapids in our data employ a strategy primarily based on TFTx, while two Micrurus species (Mc, Mf) have a combination of PLA2 and TFTx. Colubrids show a unique trend of being scattered throughout the phylomorphospace, having at least one species adopting each of the three strategies. Despite the lack of constraint in gene expression, the snake venom phenotype has very low dimensionality with the four major components accounting for 73.5% of the variation. Clustering of distantly related snakes to around a similar strategy hint at the likely parallelism of these major toxins.

The clustering of snakes on this phylomorphic venom space shows a clear association between family and the major component in the venom. For example, most elapids venoms form a cluster dominated by TFTx, which is the principal family found in their venom. On the other hand, vipers occupy a larger region of phylomorphospace because some have venoms dominated by SVMP, while others use different combinations of SVMP, SVSP and PLA2. Finally, colubrid venoms are the most diverse in composition, employing all the of the different strategies. A key observation in the PCA is that some distantly related species cluster together around the same envenomation strategy, suggesting parallel evolution.

It is important to note that PLA2s in elapids (group I) and vipers (group II) are produced by different loci and have apparently evolved independently (33, 34). In order to account for any underlying family-specific evolutionary trend, we conducted a parallel analysis by splitting PLA2 into elapid PLA2 (ePLA2) and viperid PLA2 (vPLA2) (Supplementary Fig. 5). This analysis produced qualitatively the same results as the combined analysis, though the first two components of the PCA explained less variance (61.8% as opposed to 73.6%). In particular, loadings for both elapid and viperid PLA2 were oriented in the same direction (Supplementary Fig. 10), consistent with the previous observations that they convergently evolved similar toxic activities (35). Thus, we carried out all subsequent analysis by combining them into a joint functional category.

### Parallelism of envenomation strategies

The clustering of distantly related species in the PCA despite the generally high phylogenetic inertia hinted at the likely parallelism of envenomation strategies across snakes. To test for parallelism across the phylogeny we used SURFACE (36), which fits a series of stabilizing selection models to identify instances where multiple lineages adopt the same selective regime (36). SURFACE uses AIC as criterion to determine goodness of fit, and keeps adding models until the AIC doesn’t improve further (36). The final model included 11 regime shifts and 4 distinct regimes (Δ*k* = 4) and a *c* = 7 convergent shifts. The AIC improved from 572.5 to 438.25 in the forward phase, to a final AIC of 407.56 in the backward phase (S11) which indicated that the final model was a better fit than the initial ones. The SURFACE model revealed widespread convergence in elapids, vipers, and colubrids (Fig. 3). Vipers showed evidence of three distinct regimes, out of which two evolved in parallel (Fig. 3, Supplementary 12). One of the convergent regimes focused on SVMP evolved repeatedly in viperids and colubrids (Fig. 3). Another strategy, adopted by three species across all three families (*Ovophis okinavensis, Crotalus simus*, and *Pseudonaja textilis*) was focussed around SVSP (Fig. 3). In elapids, there was greater evidence for a single convergent regime focused around TFTx. We used the inbuilt simulation function in SURFACE to obtain a null distribution on a simulated dataset using a Hansen model that lacked true convergence (36, 37). Comparison to the null model simulations (Supplementary Table 2) revealed significantly more convergent regimes obtained from our analysis than would be obtained by chance (p_c_=0.030). This allowed us to reject the null hypothesis and state that the cases of convergence are due to some optima in the phenotypic adaptive landscape.

**Fig. 3:**
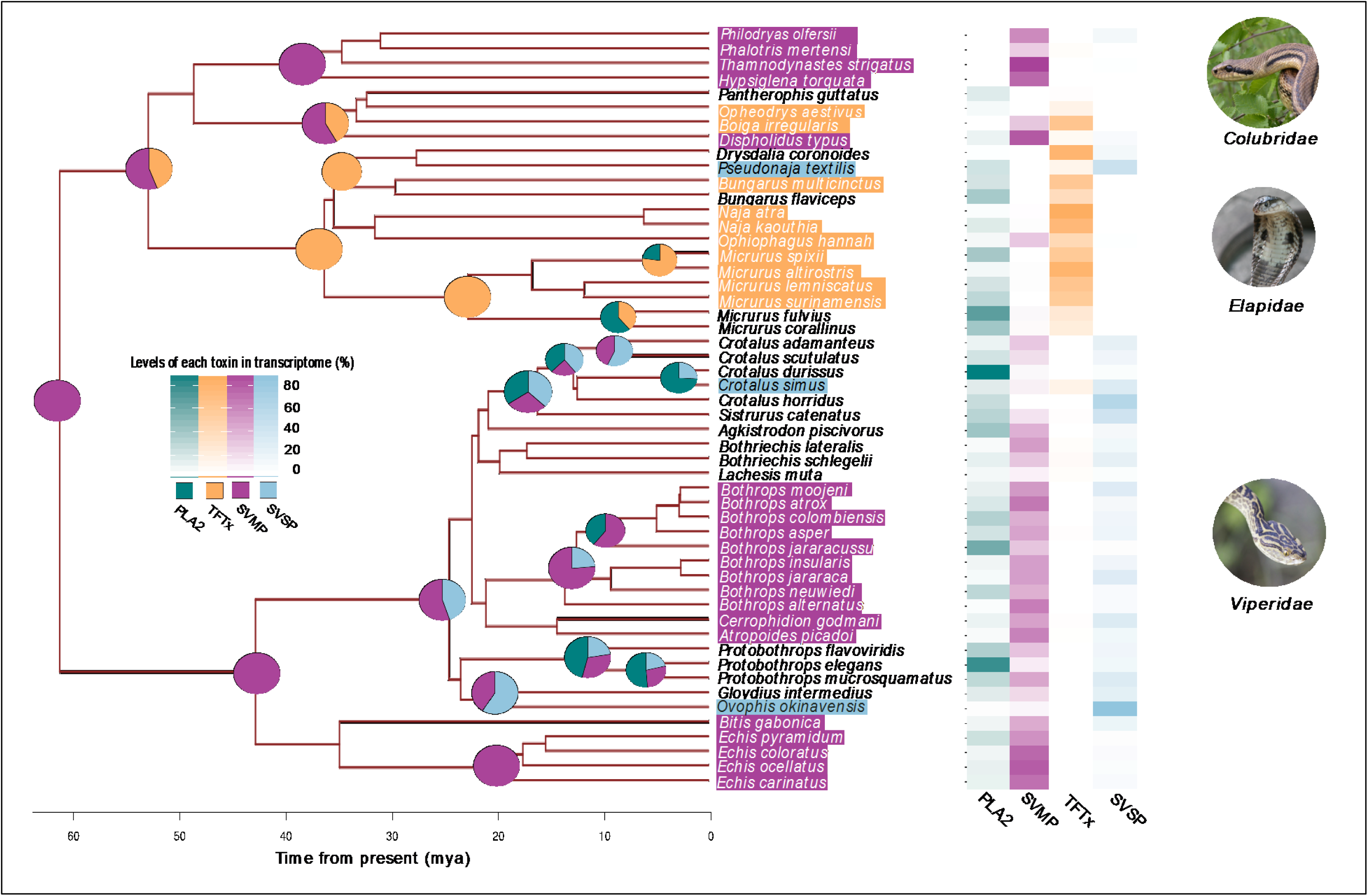
Ancestral venom (at the root 60 million years ago) was likely unspecified, and among the major components contained only SVMP. The specialization of snake venom occurred relatively recently, in the past 20-40 million years, as denoted by the ancestral state reconstruction along the nodes. Different species specialized using similar components leading to a high degree of parallelism (common selective regimes are indicated by highlighted species names). Tiles represents the relative abundance of venom toxin in extant snakes. Although ancestral states were reconstructed at each node, for clarity only the ones where substantial changes in toxin levels took place are labelled. The overall trend is that starting from an undifferentiated ancestor, snakes have increasingly focused on specific toxin families, occasionally investing into new toxin categories for their arsenals (e.g., PLA2s and SVSPs). The increased concentration of specific venom components, relative to the ancestors, has most likely happened by increases in copy number of the specific gene families.

### Strategies based on major components evolved at different times

Understanding the ancestral state of a trait can paint a picture of the journey taken by the trait through evolution. We used ancestral state reconstruction (ASR) analysis to estimate recruitment times of the major venom components into the venom arsenal, and how venoms have changed throughout the course of evolution. Because of the diversity and plasticity of the venom phenotype, confidence intervals at the root were very large, and the inference of the venom in the most recent common ancestor should be interpreted with caution, particularly concerning absence of individual toxins. Of the four major components that are responsible for venom diversification, the ASR detected only SVMP in the most common ancestor of the snakes (~60 million years ago, hence forth referred to as ‘the ancestral venom’) (Fig 3). The ASR reveals SVMP to be a major and widespread component for most of the evolutionary history of snakes. However, at the base of elapid radiation, SVMP was largely replaced by TFTx as the major component in elapid venoms. TFTx was likely present prior to the split of colubrids and elapids, but while elapids have focused primarily on TFTx, colubrids employed a combination of TFTx and SVMP throughout their evolution. In vipers SVMP has taken various paths, from being the predominant component in Viperinae (*Echis and Bitis*), to diversifying substantially in the Crotaline clade (*Protobothrops*, *Bothrops*, *Crotalus*, etc). The ASR suggests that high levels of PLA2 and SVSP (which is mostly restricted to vipers) are more recent additions to the venom. Although not shown in our analysis, PLA2 (both group I and group II) was most likely present at the common ancestors of both Elapids and Crotalids (34), but became substantial parts of the venom from around 20 million years ago in both these taxa as observed from their increased occurrence. While we had estimated ancestral states for the other 6 components as well (Supplementary Fig. 23-34), we limited our discussion to only the major toxins since they dominate adaptive optima in the venom phylomorphospace.

## Discussion

We set out to understand how changes in gene expression underlie the evolution of a complex trait, the snake venom. First, we examined the dimensionality of this trait by estimating phylogenetic covariances between expression levels of individual toxins. The covariances between toxin expression levels can be viewed as constraints that limit the evolution of a trait, analogously to the G-matrix in quantitative genetics. Unlike the G-matrix, which arises largely from pleiotropic interactions between genes, phylogenetic constraints may additionally include ecological, developmental, physiological, and other factors. Significant covariance between individual components would reflect constraints on evolutionary change and the total phenotypic space attainable by selection (38). Thus, traits that are constituted by genes under high constraint would not be able to diversify as freely as traits with no constraint. Genetic constraints also determine convergence and parallel evolution, where high constraint reduces the likelihood of genes contributing to different convergent regimes (16). Yet, for snake venom genes we see no such constraints in gene expression, suggesting that all toxin combinations, in principle, are possible (Fig. 1).

While the lack of constraint between components implies that venom has the potential to diversify freely and fully fill the possible phenotype space, this is far from what we observe. Rather, the total phenotypic space has surprisingly low dimensionality, with two principal components explaining 73% of the variance. Venoms form three distinct clusters around the major toxin components in the phylomorphospace, indicating the possible presence of three distinct adaptive optima (Fig. 2). Similar toxin-specific strategies have been observed between populations of snakes, but we show that the trend extends phylogenetically to different species as well as different families (39, 40). While individual venom components do exhibit significant phylogenetic inertia (Fig. 1), the phylomorphospace clusters often include unrelated taxa, suggesting shifts in envenomation strategies between adaptive optima. These shifts likely result from parallelism, which may be facilitated by lack of constraints between components (Fig. 3).

Is this lack of constraint surprising for a trait like snake venom? To answer this we need to understand one of the key processes by which novel functions and variations in gene families arise – gene duplication (41–44). Gene duplication can cause functional redundancy by producing gene copies where the original gene carries out its designated function while the new copy has no active role in the biological process, thus freeing it from selective constraints (41, 45, 46). This relaxed selective constraint could allow the duplicated genes to diversify freely, as long as one of the copies performs the essential function, and the presence or absence of another copy does not affect fitness. Therefore, a system that comprises of many duplicated gene families would also likely have the ability to diversify freely. Snake venom fits this characteristic since it consists of gene families that have undergone varying degrees of duplications throughout their history (47). We hypothesize that the lack of constraint observed between expression levels of genes encoding for snake venom could be due to the fact that snake venom comprises of duplicated genes.

One of the most prevalent theories about the origins of venom composition suggests that they originated after ancestral physiological genes underwent duplication and neofunctionalization (48). Since venom phenotypes need to be flexible and to adapt quickly, duplicated genes make ideal toxin candidates as they are under lower selective constraints (49–51). In addition to sequence-level changes, changes in gene expression also contribute to microevolution in snake venom (52). To get a complete picture of the evolution of the snake venom phenotype, we need to understand how microevolution (changes in gene expression over short time scales) relates to macroevolution (selection over large time scales). From our observations, we propose a model for snake venom evolution that could potentially link the two, and explain why in spite of having the potential to freely evolve, snake venom has such low dimensionality. We propose that gene duplication facilitated recruitment of physiological genes into the venom system, following which expression levels were free to respond to natural selection due to their low constraint and to potentially occupy a wide phenotypic space. The venom compositions that provided the greatest adaptive advantage due to their favourable biochemistry of envenomation is what we see in present-day species. These observed adaptive optima are dominated by the four main toxins leading to a high degree of parallelism. This model could likely explain why snake venom, like other systems comprising of duplicated genes, experience both positive and relaxed purifying selection (23, 53).

### Temporal patterns in venom evolution

Ancestral snake venom composition has received considerable attention, but until now the analyses have been qualitative in nature (39). While the confidence intervals for ancestral state reconstruction (ASR) are large (Supplementary Fig. 14-34) owing to the remarkable evolutionary lability of venom, we can nonetheless make a number of observations about the course of evolution of major components. Among the major components, the ancestral venom most likely contained only appreciable amount of SVMP (Fig. 3). This finding is consistent with previous estimates of a likely recruitment of SVMP into the venom at the split of vipers and colubrids (~62 million years ago) (24, 54). Furthermore, the SVMP-focused strategy is the only convergent selective regime identified by the SURFACE analysis in all three families (Fig. 3), suggesting that the machinery to produce this toxin exists in all of them. While we could not detect PLA2, TFTx, and SVSP with confidence in the most recent common ancestor, they could have been present at lower levels in the ancestral venom, or as ancestral precursor molecules (33, 34, 55). This is especially likely given that all three families have some level of each of the major toxin classes (Fig. 3).

Being present in the ancestral venom, SVMP continued to be used as a major toxin by viperids and is still the dominant toxin in some genera (*Echis and Bitis*), as well as some colubrids. However, other toxins were recruited (or increased in quantity) later in venomous snake evolution. For example, consistent with previous work that placed recruitment of TFTx before the divergence of modern elapids (56), we also show that TFTx was likely present prior to the split between elapids and colubrids. At that time TFTx may have co-occurred with SVMP prior to the split of Elapids and Colubrids, perhaps as a specific strategy, one that is quite rare in present-day snakes, being found only in the colubrid brown tree snake (*Boiga irregularis*), and to an extent in the king cobra (*Ophiophagus hannah).* With the proliferation of the TFTx family elapids have largely lost their reliance on SVMPs.

Viperid and elapid sub-families have convergently evolved greater reliance on PLA2 toxins (group I in elapids and group II in viperids), but have diverged in venom phenospace due to the previous co-option of different major components (TFTx for elapids, SVSP for vipers). The likely presence of PLA2 (group II) gene copies at the common ancestors of Crotalids raises questions about when the complex expanded in the course of snake evolution (34). From our analysis, we believe that the expansion started somewhere around 20-25 million years ago in vipers, and was already established as a substantial part of the venom before the split of *Crotalus*, and *Protobothrops* genera. In elapids ASR does not detect the use of PLA2 before its recruitment as a major component of coral snakes (Micrurus) about 20 million years ago, but it was likely present at the common ancestor of elapids and maybe even colubrids because of its presence in many extant species. Interestingly, the recruitment of the two PLA2 families by elapids and viperids occurred at roughly the same time, perhaps as a result of convergent selection driven by radiations in prey lineages, such as mammals.

The overall trend is that recruitment of major toxins took place at different times, and has progressed along different trajectories in different lineages, with instances of both loss and heightened expression. Snakes have then shifted focus on specific toxin families, occasionally investing into new toxin categories for their arsenals (e.g., PLA2s and SVSPs). The increased concentration of specific venom components, relative to the ancestors, has most likely happened by increases in copy number of the specific gene families (47, 48, 52). Interestingly, shifts in selective regimes produced parallel specialization on the same toxin family by different snakes (Fig. 3), suggesting that at the level of toxin family selection generally favours specialization as opposed to diversity.

## Conclusion

The extent to which traits are constrained by their history, vs reaching their fitness optima has been a major debate in evolutionary biology. Numerous studies have relied on phylogenetic regression to estimate morphological covariation between traits while accounting for phylogenetic non-independence (20, 57–60). In our approach we analyse more than one response variable simultaneously and incorporate effects on trait relationships that arise through shared ancestry (61). We show that the structure of the gene expression PCOV can give insights into how traits evolve, by providing a conceptual bridge between micro and macroevolutionary forces. By showing that the phenotypic space is inherently unconstrained, we are able to highlight the existence of fitness optima, and explain the existence of widespread parallelism seen in snake venoms. These findings show that in the long-term snakes are able to overcome the inherent trade-off between fitness and phylogenetic constraints. Once genes underlying more traits are known in other systems, subsequent studies will show to what extent snake venoms are typical of the general evolutionary pattern.

## Materials and Methods

### Data collection

Toxin expression data was collected from 39 publications (online supp), while mean size measurements were obtained from Encyclopedia of life and The Reptile Database (62, 63). Out of the vast repertoire of venom toxins we selected only 10 as they were the most reported toxins amongst all snakes. Toxins levels were recorded as per publication. Toxin values reported as absolute FPKM values, were converted to a percentage of the total. All analyses were carried out using this curated dataset. The toxin values were normalised for calculating the PCOV and in SURFACE analyses. Measurements of snake size (total length, average length, snout-vent length) as reported in the online databases (62, 63) was used in the analysis. If the length was reported as a range, midpoint value was recorded in the dataset.

### Phylogenetic tree

We used a time-calibrated tree of squamate reptiles (snakes and lizards) based on two large datasets comprising of 44 nuclear genes for 161 squamates, and a dataset of 12 genes from 4161 squamate species, both these datasets represented families and subfamilies (27, 64, 65). The result was an extensive phylogeny of squamates both in terms of sampling of genes and species. Fossil based age constraints were used in time-calibrating the tree making it ideal for studies of biogeography, diversification and trait evolution (27). All analysis was carried out using a pruned version of this tree (Supplementary Fig. 14) that contained the 52- snake species for which we collected gene expression data. This pruned tree had a time at root estimated to be approximately 60 million years ago.

### Estimating Phylogenetic covariance matrix

The effect of phylogeny was modelled using the method stated in section 11.2.1- “General Quantitative Genetic Methods for Comparative Biology” (29). Analysis was carried using the MCMCglmm package in R (61). The model was written based on the description given in section 3 on the MCMCglmm vignette for modelling multi-response traits (61). Phylogenetic generalised linear mixed models allow for testing slightly complicated models, provide more than a simple qualitative estimate of the existence of phylogenetic structure, and have greater statistical power than typically used metric randomization approaches (66). The MCMC was run for a total of 20 million iterations, with burnin and thinning values of 500,000 and 1,500 respectively. Diagnostics for the MCMC run were done by obtaining the plot for the MCMC and autocorrelation. The phylogenetic signal was obtained by dividing the covariance for each toxin by the total covariance of the toxin and the residuals, as mentioned in (29). We performed principal components analysis using the phylogenetic covariances obtained from the MCMCglmm analysis. Species codes are provided in supplementary note 1.

### Convergence analysis

We used the default Ornstein-Uhlenbeck process, a convenient representation of evolution towards adaptive peaks for modelling convergence in the SURFACE analysis (36). The SURFACE method uses Hansen’s approach (Hansen model) of modelling evolution towards different adaptive optima by painting multiple adaptive hypothesis onto branches of a phylogenetic tree(36, 67). SURFACE is unique because unlike previous methods that utilize Hansen models, the placements of regime shifts is guided by trait data as opposed to some a priori hypothesis regarding the location of convergence (36). The SURFACE method is divided into two phases. The forwards phase adds successive regimes to a basic Hansen model using input from continuous trait measurements, which in our study were normalized measurements of gene expression for the four major toxins. The performance of each successive model was measured using AIC by balancing improvements in log-likelihood against increase in model complexity (36). Since AIC for the models are calculated after adding log-likelihoods, the AIC for successive models may improve. The regime shift representing the best model is painted onto the tree. The backwards phase is the second phase in the analysis. During this phase of SURFACE all subsets of regimes are collapsed to yield distinct regimes. The collapse is continued till the AIC of the models does not increase further. The final model has *k* regime shifts, and *k*’ distinct regimes, in addition to the extent of convergence which is defined as the difference of these terms (Δ*k*), *c* is used to represent the the shifts towards different convergent regimes in multiple lineages (36). We used all standard parameters as mentioned in the SURFACE vignette (37). To obtain a null distribution we ran 500 iterations of the in-built *surfaceSimulate* function using a Hansen-fit model, and concatenated the output from each iteration.

### Ancestral state reconstruction (ASR)

The default parameters for the *fastAnc* function implemented in the Phytools package was used to perform the ASR (68). A phenogram, which shows relative positions of species in evolutionary phenospace, was plotted for each toxin using a spread cost of 0.1 (Supplementary Fig. 15-34). We used the contMap function in Phytools to obtain a tree for changing trait values on a continuous scale represented by a color spectrum. Confidence intervals were plotted on the nodes as bars. Only traits whose confidence intervals did not overlap zero (only positive values) were considered to be present at the root. Pie charts in the main figure were drawn by calculating the relative levels of each of the major toxins estimated by the ASR at the specific node. Two images in the main were obtained from Wikimedia under the creative commons license (Elapidiae: Thomas Jaehnel, Colubridae: Carlo Catoni) image for Viperidae provided by Alexander S. Mikheyev)

## Acknowledgements

We would like to thank all the members of the Ecology and Evolution unit at OIST for their input and feedback. Ivan Koludarov and Steven D Aird for useful discussions about snake venom biochemistry and evolution. We are especially grateful to Steven Aird for locating several additional data sets. Nick Friedman form the Biodiversity and Biocomplexity unit at OIST for discussions regarding comparative methods.

## Author contributions

Dataset was collected by AB. Both AB and ASM analysed the data. AB and ASM wrote the paper.

## Additional information

Supplementary information, including code, data, original figures are available at: https://agneeshbarua.github.io/Many-options-supplementary/

